# Protease-Activated Receptor 2 (PAR2) expressed in sensory neurons contributes to signs of pain and neuropathy in paclitaxel treated mice

**DOI:** 10.1101/2023.02.12.528175

**Authors:** Moeno Kume, Ayesha Ahmad, Kathryn A. DeFea, Josef Vagner, Gregory Dussor, Scott Boitano, Theodore J. Price

**Author notes:** **Corresponding Author:** Theodore Price, Ph.D.; Professor, Department of Neuroscience and Center for Advanced Pain Studies; University of Texas at Dallas; 800 N. Campbell Rd; Richardson, TX 75080; (972)883-4311. **Author Contributions:** MK, GD, JV, TJP, SB, and KAD all conceived and designed the project. MK, GD, JV, TJP, KAD, and SB all regularly discussed and refined the project. MK and AA performed experiments and collected data. MK and TJP analyzed data. MK and TJP wrote the first draft of the paper. All authors read, contributed with writing and/or edited the final manuscript.

## Abstract

**Background and Purpose:** Chemotherapy-Induced Peripheral Neuropathy (CIPN) is a common, dose-limiting side effect of cancer therapy. Protease-activated receptor 2 (PAR2) is implicated in a variety of pathologies, including CIPN. In this study, we demonstrate the role of PAR2 expressed in sensory neurons in a paclitaxel (PTX)-induced model of CIPN in mice.

**Experimental Approach:** CIPN was induced in both PAR2 knockout/WT mice and mice with PAR2 ablated in sensory neurons via the intraperitoneal injection of paclitaxel. *In vivo* behavioral studies were done in mice using von Frey filaments and the Mouse Grimace Scale. We then examined immunohistochemical staining of dorsal root ganglion (DRG) and hind paw skin samples from CIPN mice to measure satellite cell gliosis and intra-epidermal nerve fiber (IENF) density. Pharmacological reversal of CIPN pain was tested with the PAR2 antagonist C781

**Key Results:** Mechanical allodynia caused by paclitaxel treatment was alleviated in PAR2 knockout mice of both sexes. In the PAR2 sensory neuronal conditional knockout (cKO) mice, both mechanical allodynia and facial grimacing were attenuated in mice of both sexes. In the dorsal root ganglion of the paclitaxel-treated PAR2 cKO mice, satellite glial cell activation was reduced compared to control mice. IENF density analysis of the skin showed that the paclitaxel-treated control mice have a reduction in nerve fiber density while the PAR2 cKO mice had a comparable skin innervation as the vehicle-treated animals. Similar results were seen with satellite cell gliosis in the DRG where gliosis induced by PTX was absent in PAR cKO mice. Finally, C781 was able to transiently reverse established PTX-evoked mechanical allodynia.

**Conclusions and Implications:** Our work demonstrates that PAR2 expressed in sensory neurons plays a key role in paclitaxel-induced mechanical allodynia, spontaneous pain and signs of neuropathy, suggesting PAR2 as a possible therapeutic target in multiple aspects of paclitaxel CIPN.

## INTRODUCTION

More than 16.9 million cancer survivors were alive in 2019, and due to better early detection of cancer and advancements in chemotherapeutic agents, this number is projected to increase to more than 22.1 million by 2030 ^17, 40, 64^. With increased cancer survivorship, a common consequence of treatment that cancer patients and survivors deal with are the long-lasting neuropathic symptoms from peripheral nerve damage caused by neurotoxic chemotherapeutics, a condition known as chemotherapy-induced peripheral neuropathy (CIPN) ^27, 28, 65, 76, 85^. CIPN affects over 60% of patients in the first month after chemotherapy and, although the prevalence of CIPN decreases over time, 30% of patients continue to suffer from CIPN at 6 months or more ^79^. Common symptoms of CIPN include numbness, tingling, hyperalgesia, and spontaneous pain which can be disabling and impair quality of life ^23, 56, 70, 79, 85, 88^. Furthermore, CIPN symptoms can lead to dose reduction or even treatment cessation, potentially affecting patient survival ^8, 23, 70, 82^. Despite this, few effective therapeutics are currently available to remedy or prevent the effects of CIPN due to a poor understanding of the underlying mechanism ^24, 33, 59^.

The effects of chemotherapy drugs on the nervous system can vary depending on the dosage and drug class, the most neurotoxic classes being platinum-based drugs, taxanes, and thalidomide ^6^. Paclitaxel (PTX) is a taxane chemotherapeutic drug used in the treatment of breast, lung, and ovarian cancer, among others ^93, 96^. Taxanes impair cancer cell division and growth by stabilizing microtubules, thereby preventing the chromosomes from separating and leading to apoptosis in dividing cells ^1, 5, 37, 99^. Though neurons are not dividing cells, they are still susceptible to complications from PTX exposure. PTX has poor penetration into the CNS but has been shown to accumulate in the DRG at high concentrations, thus resulting in mainly sensory neuropathic symptoms ^13, 21^. Effects on the PNS by PTX are multifaceted and include dysfunction of microtubules, mitochondria, and calcium homeostasis, axon degeneration, hyperexcitability of sensory neurons, and activation of glial and immune cells ^76, 99^.

Protease-Activated Receptor 2 (PAR2) has been proposed as a possible mediator of CIPN pain via the neuroimmune interactions that are catalyzed by chemotherapy treatment ^15, 75, 86^. PAR2 is a G-protein coupled receptor (GPCR) implicated in a number of pathologies including inflammatory, cancer, migraine, and CIPN pain ^15, 30, 46, 49, 50, 61, 74, 75, 86, 90, 101^. The extracellular portion of the receptor is cleaved by proteases, such as those released by immune cells. Cleavage by trypsin-like proteases exposes a tethered ligand that can then bind to the receptor, leading to the recruitment of G_q_ and β-arrestins, while cleavage by elastase promotes the recruitment of G_12/13_ and Rho-dependent activation of MAPK ^2, 34, 42, 47, 73^. The β-arrestin/MAPK pathway in particular causes nascent protein synthesis in a small subpopulation of sensory neurons and causes mechanical hypersensitivity and facial grimacing after PAR2 activation ^32, 87^.

Here, we test the hypothesis that sensory neuronal PAR2 plays a key role in CIPN pain using behavioral genetics, pharmacology and immunohistochemistry. We measured both mechanical and spontaneous nociceptive behaviors using von Frey filaments and the Mouse Grimace Scale (MGS), respectively, in paclitaxel-treated PAR2 global knockout mice and *Pirt*^*Cre*^*F2rl1*^*flox*^ mice. We also measured nerve fiber density in the hind paw skin and satellite cell gliosis in the dorsal root ganglion (DRG) in the *Pirt*^*Cre*^*F2rl1*^*flox*^ mice after paclitaxel treatment. Finally, we tested a PAR2 antagonist, C781, in a paclitaxel-induced CIPN model. Our findings demonstrate that PAR2 expressed on sensory neurons contributes to both mechanical and spontaneous nociceptive behaviors as well as changes in nerve fiber density in the epidermis and satellite cell gliosis.

## MATERIALS AND METHODS

### Animals

All experiments and procedures were approved by the Institutional Animal Care and Use Committee at University of Texas at Dallas. To generate *Pirt*^*Cre*^*F2rl1*^*flox*^ mice, *loxP* sites were inserted on exon 2 of the *F2rl1* gene and crossed with the *Pirt*^*Cre*^ mouse, as described previously ^32^. Our experimental mice were homozygous for the *F2rl1*^*flox*^ gene and heterozygous for the *Pirt*^*Cre*^ gene due to the insertion of Cre recombinase into the *Pirt* promoter, thus rendering Pirt expression null in the allele. For control groups, we used vehicle-treated mice as well as *Pirt*^*Cre*^ mice that have wildtype *F2rl1* expression. The mice were bred in house at the University of Texas at Dallas and housed in climate-controlled rooms set to 22ºC with a 12-hour light/dark cycle and given food and water *ad libitum*. Male and female mice were used in all experiments. Because we have not previously found any sex differences in PAR2 signaling related to pain, sex differences were not assessed in experiments described here and the study was not powered to evaluate sex differences.

### Experimental reagents

Paclitaxel was purchased from Sigma Aldrich (Y0000698) and dissolved in a 50/50 Kolliphor® EL (Sigma Aldrich C5135)/ethanol solution and then further diluted in sterile Dulbecco’s phosphate buffered saline (DPBS; Thermo Scientific) for injections. Paclitaxel was injected into the peritoneal space using a 27G ½” needle. C781 was synthesized using solid-phase synthesis as described previously ^78^. C781 was injected into the peritoneal space using a 27G ½” needle at a dosage of 10 mg/kg.

### Behavioral Methods

Experimenters were blinded to treatment and genotype groups in all experiments. CIPN was induced by injecting PTX (4 mg/kg IP) every other day for a total of four injections (cumulative dose of 16 mg/kg) ^39, 62, 89^. Intermittent systemic administration of PTX was done to mimic cyclic chemotherapy dosing regimens in human patients ^25, 36^. Animals were habituated to suspended acrylic chambers with each individual chamber measuring 11.4 × 7.6 × 7.6 cm and a wire mesh bottom (1 cm^2^). Mechanical hypersensitivity was assessed using the manual von Frey hair test after spontaneous pain assessment using the Mouse Grimace Scale (MGS). This was done to assure that grimacing responses were not a result of mechanical stimulation.

Mouse grimace scores were determined using the Mouse Grimace Scale (MGS) test as described by Langford and colleagues ^52^. After habituation, the experimenter observed the mice and assigned intensity ratings (0 = not present, 1 = moderately present, 2 = severe) to the facial action units relative to baseline. The five action units identified are 1) orbital tightening, 2) nose bulge, 3) cheek bulge, 4) ear position, and 5) whisker change. The scores for all five action units are averaged to obtain the MGS score for each time point. The MGS has been shown to be an accurate way of measuring spontaneous acute nociception in response to multiple types of stimuli, including PTX-induced CIPN ^3, 4, 54, 60, 91^.

Hind paw withdrawal thresholds were calculated via the Dixon up-down method after applying calibrated von Frey filaments from Stoelting, Co. (cat# 58011) to the volar surface of the left hind paw ^14^. Increasing pressure was applied until the filament bent against the skin at a right angle and a positive response was recorded when the mouse showed nocifensive behaviors, such as rapid paw withdrawal, flicking, and/or licking of the stimulated paw. The maximum filament weight used was 2 g.

### Immunohistochemistry

After the conclusion of behavioral experiments, animals were anesthetized using 4% isoflurane and euthanized by cervical dislocation and decapitation. Hind paw skin was carefully dissected and fixed overnight in Zamboni’s fixative solution at 4°C. The skin was washed three times (30 min each) in PBS to remove excess fixative and incubated in 30% sucrose at 4°C until it sunk. Samples were blotted dry on paper towels, embedded in Optimum cutting temperature (OCT), and flash-frozen in dry ice. Tissues were sectioned at 50 µm and immersed into a netted well with PBS. Sections were washed three times in PBS to remove excess OCT and then incubated in blocking buffer (10% normal goat serum, 0.3% Triton X-100 in 0.1 M PB) for 1 hr at room temperature. Sections were then incubated overnight in primary antibody (rabbit anti-PGP9.5; CL7756AP-50; Cedarlane) at 1:1000. The next day, sections were washed in PBS three times and incubated in secondary antibody, cross-adsorbed goat anti-rabbit IgG (H+L) Alexa Fluor 555 (1:2000; A21428; Invitrogen, Thermo Fisher Scientific) for 1 hr at room temperature. Finally, sections were incubated for 5 min in 1:5000 DAPI (ACD), washed in PBS two times, moved and flattened onto slides, and cover-slipped using ProLong Gold mounting medium (Thermo Fisher Scientific).

Dorsal root ganglia (DRGs) were dissected and embedded in OCT and immediately flash-frozen in dry ice. Tissues were sectioned at 20 µm onto charged slides where they were briefly thawed to adhere to the slide but immediately returned to the -20°C cryostat chamber until completion of sectioning. The sections were fixed in ice-cold 10% formalin for 15 min followed by incubation in 50%, 70%, and then 100% ethanol for 5 min each. Slides were placed in a light-protected, humidity-controlled tray and incubated in blocking buffer for 1 hr at room temperature. Slides were then incubated overnight in mouse anti-GFAP (N206A/8; NeuroMab) and chicken anti-peripherin (CPCA-Peri; Encor Biotechnology) at 1:1000 in blocking buffer overnight at 4°C. The next day, slides were washed in PBS three times and incubated in goat anti-chicken (H&L) Alexa Fluor 488 (A11039; Invitrogen, Thermo Fisher Scientific) goat anti-mouse IgG1 Alexa Fluor 555 (A21127; Invitrogen, Thermo Fisher Scientific) at 1:2000 for 1 hr at room temperature. Finally, sections were incubated for 5 min in 1:5000 DAPI, washed in PBS two times, and then cover-slipped using ProLong Gold mounting medium.

### Image analysis for skin sections

Three sections per mouse were imaged on an Olympus FV3000 confocal microscope at magnification x20 and then analyzed using Cellsens (Olympus) software. Acquisition parameters were maintained constant. To determine the number of intraepidermal nerve fibers (IENF), the number of free nerve endings that cross the dermo-epidermal junction was counted in a blinded manner according to counting rules recommended by the European Federation of Neurological Societies (EFNS) ^53^. The IENF density was calculated as the number of fibers per millimeter of epidermal length.

### Image analysis for DRG sections

Three sections per mouse were imaged on an Olympus FV3000 confocal microscope at magnification x20 and then analyzed using Cellsens (Olympus) software. Acquisition parameters were maintained constant. Satellite glial cells were distinguished via GFAP staining, as well as shape and position in relation to neurons, which were stained with peripherin. Neurons that were surrounded by satellite glial cells by half or more of their circumference were categorized as GFAP^+^ neurons. The total number of GFAP^+^ neurons were quantified and divided by the total number of neurons in the field and presented as a percentage.

### Data and Statistical Analysis

Sample sizes for *in vivo* experiments were estimated based on effect sizes observed in PAR2 sensory neuron-specific conditional knockout (cKO) mice as reported previously ^32^. All statistical tests were done using GraphPad Prism version 9.4.1 (GraphPad Software, Inc.). We assessed whether the data violated assumptions of parametric statistics via the Brown-Forsythe test and the Kolmogorov-Smirnov test when doing the analysis in GraphPad Prism. If the standard deviations were not significantly different and the data passed the normality test (alpha > 0.05), differences between groups were calculated using repeated measures 2-way ANOVAs with Dunnett’s or Sidak’s multiple-comparison test. Effect size was determined by calculating the cumulative differences between the baseline value and the experimental value at each time point. Two-way ANOVA with Dunnett post-test was used to compare effect sizes. Statistical analysis for IENF and SGC analysis is described in figure legends. All data are represented as mean ± SEM.

## RESULTS

### Mechanical hypersensitivity and grimace caused by PTX treatment are profoundly reduced in mice lacking PAR2 expressed on sensory neurons

We tested whether global PAR2 knockout mice exhibit reduced mechanical hypersensitivity in a paclitaxel-induced CIPN model. To induce CIPN, we injected 4 mg/kg of PTX into the intraperitoneal space every other day for a total of 4 injections. Using von Frey testing, we found that PTX caused long-lasting mechanical hypersensitivity in the wildtype control mice that lasted out to 23 days. In contrast, this mechanical hypersensitivity was significantly attenuated in the global PAR2 knockout mice at day 9, 11, 14, 19, and 23 **(Figure 1A)**. The vehicle-treated control animals showed paw withdrawal thresholds similar to baseline levels. The WT PTX mice showed significantly higher effect size when compared to both vehicle groups but not to the PTX-treated PAR2 knockout group **(Figure 1B)**.

**Figure 1:**
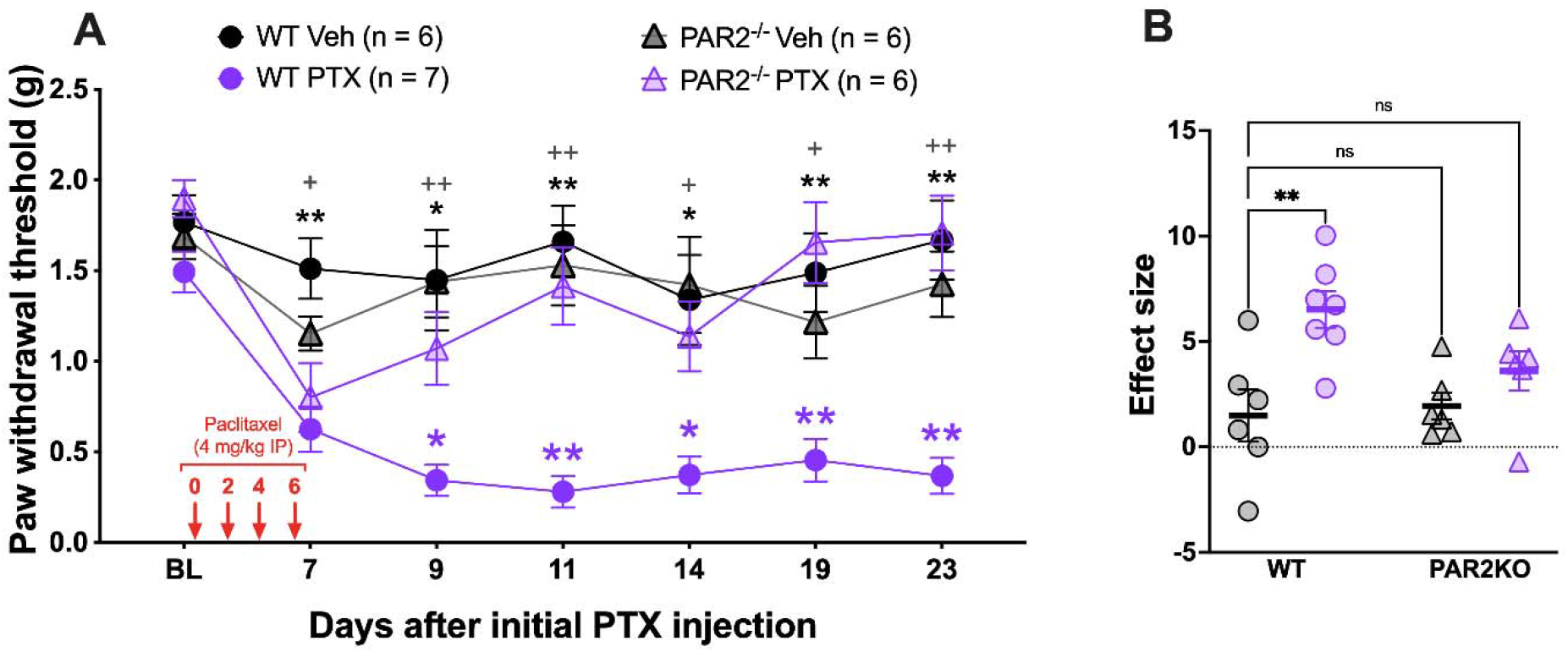
PAR2^-/-^ mice show attenuated mechanical hypersensitivity in response to paclitaxel treatment. Baseline (BL) paw withdrawal thresholds for male and female ICR mice were obtained before paclitaxel treatment (4 mg/kg IP every other day for 4 days). **(A)** The WT control mice that received PTX treatment exhibited long-lasting mechanical allodynia. On the other hand, the PAR2^-/-^ mice treated with PTX show significantly higher mechanical thresholds at day 9, 11, 14, 19, and 23 after the initial PTX injection. Purple asterisks denote significant differences between the WT PTX group and the PAR2^-/-^ PTX group. Black asterisks denote significant differences between the WT PTX group and the WT Veh group. Gray plus signs denote significant differences between WT PTX group and the PAR2^-/-^ Veh group. **(B)** Effect size was determined by calculating the cumulative differences between the baseline value and the experimental value at each time point. The WT PTX mice show a significantly higher effect size when compared to the WT Veh group. However, both the effect sizes of the PAR2^-/-^ groups were not significantly different from the WT Veh group. For the WT Veh group, n = 3 females and n = 3 males. For the WT PTX group, n = 4 females and n = 3 males. For the PAR2^-/-^ Veh group, n = 3 females and n = 3 males. For the PAR2^-/-^ PTX group, n = 3 females and n = 3 males. *p <0.05, **p<0.01. For paw withdrawal threshold, repeated measures two-way ANOVA and Dunnett’s multiple comparisons test was used with group mean comparisons made to the WT PTX group. For effect size, two-way ANOVA with Dunnett’s multiple comparisons test was use with group mean comparisons made to the WT Veh group.

Our previous research on PAR2-evoked pain using the *Pirt*^*Cre*^*F2rl1*^*flox*^ mouse, a sensory neuron-specific cKO PAR2 mouse, revealed that mechanical hypersensitivity and spontaneous pain caused by PAR2 activation is dependent on PAR2 expressed on a small subpopulation of sensory neurons ^32, 43, 44^. Therefore, we induced CIPN in these mice to determine whether we would we see similar results after PTX treatment. As previously seen, PTX caused long-lasting mechanical hypersensitivity in the *Pirt*^*Cre*^*F2rl1*^*+/+*^ control animals that lasted out to 23 days **(Figure 2A)**. PTX also caused facial grimacing in these animals that lasted out to 17 days **(Figure 2C)**. Compared to the *Pirt*^*Cre*^*F2rl1*^*+/+*^ control animals, the *Pirt*^*Cre*^*F2rl1*^*flox*^ mice showed similar withdrawal thresholds and facial grimace scores to the vehicle-treated animals. For von Frey testing, the *Pirt*^*Cre*^*F2rl1*^*flox*^ mice were significantly different from the *Pirt*^*Cre*^*F2rl1*^*+/+*^ control animals at day 7, 9, 13, and 17 and through overall effect size **(Figure 2A and 2B)**. For facial grimace scoring, the *Pirt*^*Cre*^*F2rl1*^*flox*^ mice were significantly different from the *Pirt*^*Cre*^*F2rl1*^*+/+*^ control animals at day 3, 7, 9, 13, and 17 and through overall effect size **(Figure 2C and 2D)**. These results suggest that the mechanical hypersensitivity and facial grimacing caused by PTX can be mediated by PAR2 expressed on sensory neurons.

**Figure 2:**
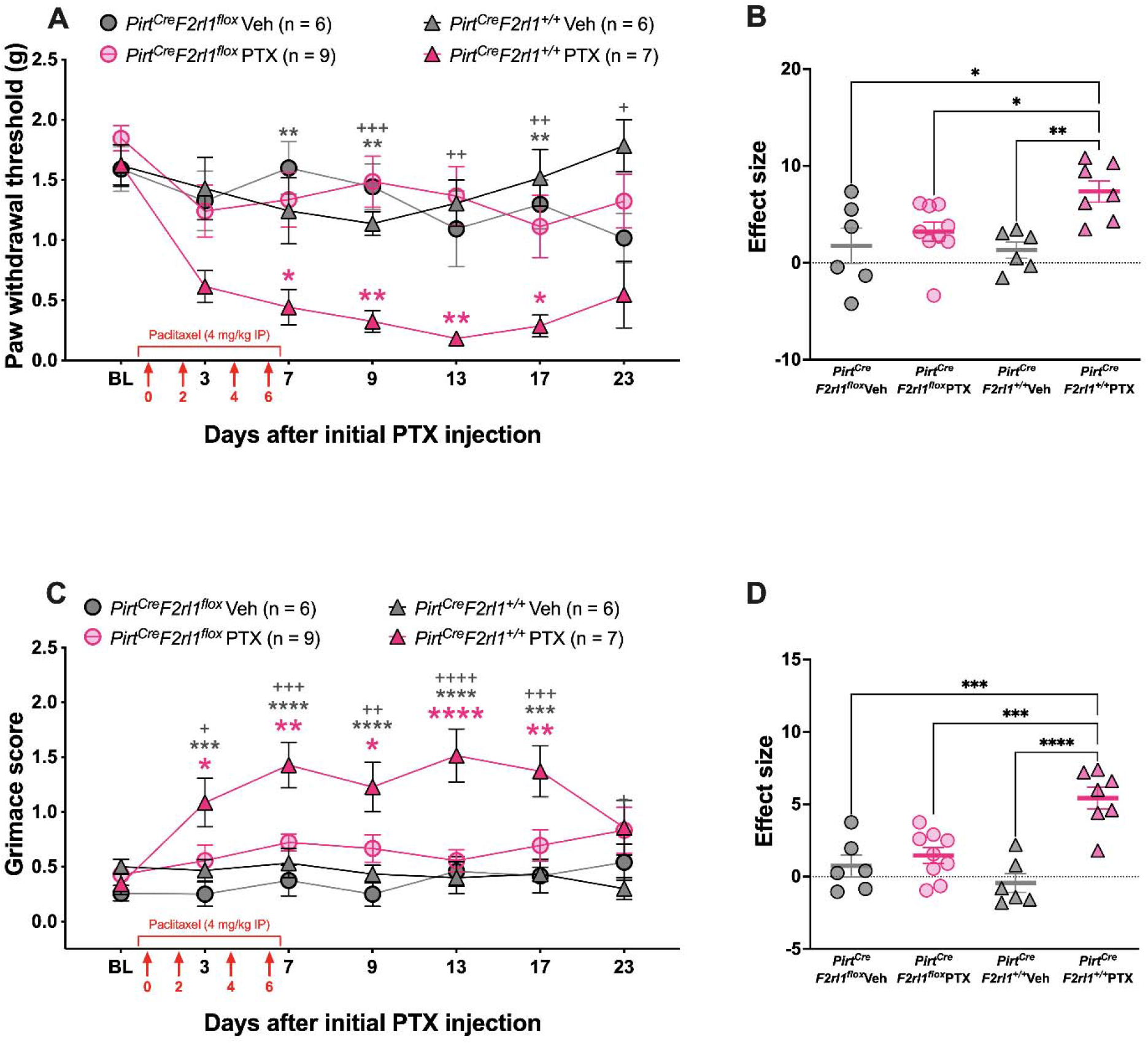
*Pirt*^*Cre*^*F2rl1*^*flox*^ mice show attenuated mechanical hypersensitivity and facial grimacing in response to paclitaxel treatment. Baseline (BL) paw withdrawal thresholds for male and female *Pirt*^*Cre*^*F2rl1*^*flox*^ and control mice were obtained before paclitaxel treatment (4 mg/kg IP every other day for 4 days). **(A)** The control mice that received PTX treatment exhibited long-lasting mechanical allodynia. On the other hand, the *Pirt*^*Cre*^*F2rl1*^*flox*^ mice treated with PTX show significantly higher mechanical thresholds at day 7, 9, 13, and 17 after the initial PTX injection. Pink asterisks denote significant differences between the *Pirt*^*Cre*^*F2rl1*^*+/+*^ PTX group and the *Pirt*^*Cre*^*F2rl1*^*flox*^ PTX group. Gray asterisks denote significant differences between the *Pirt*^*Cre*^*F2rl1*^*+/+*^ PTX group and the *Pirt*^*Cre*^*F2rl1*^*flox*^ Veh group. Gray plus signs denote significant differences between *Pirt*^*Cre*^*F2rl1*^*+/+*^ PTX group and the *Pirt*^*Cre*^*F2rl1*^*+/+*^ Veh group. **(B)** Effect size was determined by calculating the cumulative differences between the baseline value and the experimental value at each time point. The *Pirt*^*Cre*^*F2rl1*^*+/+*^ PTX mice show a significantly higher effect size when compared to all other groups. **(C)** The control mice that received PTX treatment also exhibited a long-lasting facial grimacing. *Pirt*^*Cre*^*F2rl1*^*flox*^ mice treated with PTX show significantly lower facial grimace scores at day 3, 7, 9, 13, and 17 after the initial PTX injection. **(D)** The *Pirt*^*Cre*^*F2rl1*^*+/+*^ PTX mice show a significantly higher effect size when compared to all other groups. For the *Pirt*^*Cre*^*F2rl1*^*+/+*^ Veh group, n = 3 females and n = 3 males. For the *Pirt*^*Cre*^*F2rl1*^*+/+*^ PTX group, n = 4 females and n = 3 males. For the *Pirt*^*Cre*^*F2rl1*^*flox*^ Veh group, n = 2 females and n = 4 males. For the *Pirt*^*Cre*^*F2rl1*^*flox*^ PTX group, n = 3 females and n = 6 males. *p <0.05, **p<0.01, ***p<0.001, ****p<0.0001. For paw withdrawal threshold, repeated measures two-way ANOVA and Dunnett’s multiple comparisons test was used. For facial grimace scores, repeated measures two-way ANOVA with Dunnett’s multiple comparisons test was used. For effect size, two-way ANOVA with Dunnett’s multiple comparisons test was used. All group mean comparisons were made to the *Pirt*^*Cre*^*F2rl1*^*+/+*^ PTX group.

### Epidermal innervation loss caused by PTX treatment is reduced in mice lacking PAR2 expressed on sensory neurons

PTX treatment has been demonstrated to result in a loss of IENF density in both humans and rodents ^9, 16, 45, 57, 80, 97^. Our previous analysis of single-cell RNA-sequencing and RNAscope validation experiments showed that the PAR2 gene, *F2rl1*, is expressed specifically in a small subpopulation of neurons that also express *IL31ra, Nppb, Hrh1*, and *Mrgprx1*, all genes believed to be important for mediating itch and expressed in sensory neurons that innervate the epidermis ^32, 51, 55, 63, 66^. To determine whether PAR2 expression on this subpopulation of skin-innervating sensory neurons is critical for PTX-induced IENF loss, we dissected the hind paw skin from the mice used in the *in vivo* behavioral assays after completion of experiments. The skin samples were stained for PGP9.5 to visualize the intraepidermal nerve fibers and the free nerve endings that cross the dermo-epidermal junction were counted.

The vehicle-treated animals showed dense nerve fibers that innervate the epidermis. After PTX treatment, the *Pirt*^*Cre*^*F2rl1*^*+/+*^ control mice show a clear loss of nerve fibers in the skin that is not seen in the *Pirt*^*Cre*^*F2rl1*^*flox*^ PTX group **(Figure 3A & 3B)**. The IENF density was calculated as the number of fibers that crossed the dermo-epidermal junction per millimeter of epidermal length. The *Pirt*^*Cre*^*F2rl1*^*+/+*^ PTX group showed significantly lower IENF/mm than the *Pirt*^*Cre*^*F2rl1*^*flox*^ PTX group and the *Pirt*^*Cre*^*F2rl1*^*+/+*^ Veh group. These results indicate that the *Pirt*^*Cre*^*F2rl1*^*flox*^ mice are protected from nerve fiber loss as a result of PTX treatment.

**Figure 3:**
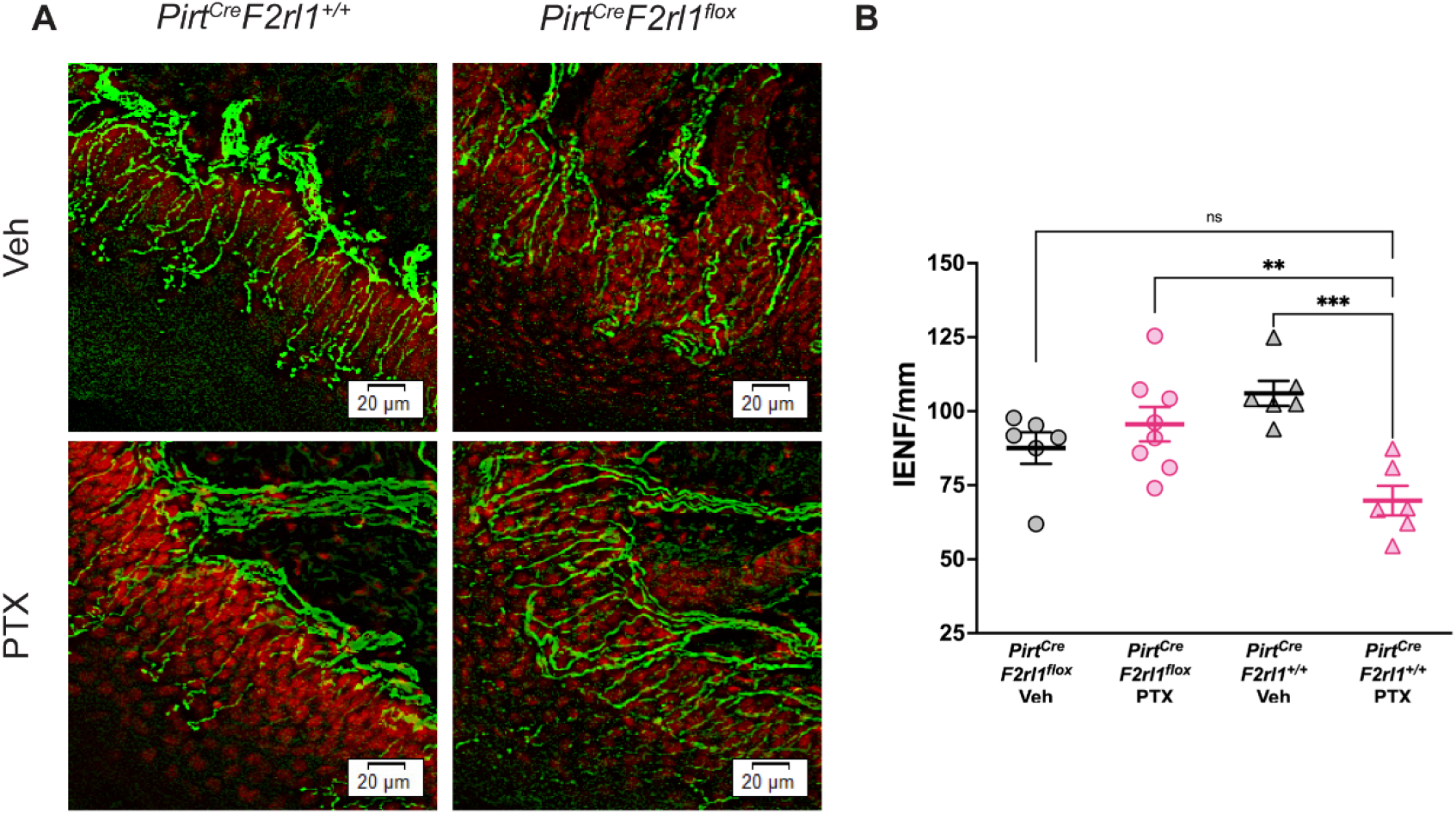
*Pirt*^*Cre*^*F2rl1*^*flox*^ mice are protected from nerve fiber loss as a result of paclitaxel treatment. **(A)** Hind paw skin was dissected, cryosectioned at 50 µm, and stained for PGP9.5 (green) to visualize skin nerve fibers and DAPI (red) to visualize the dermo-epidermal junction. The *Pirt*^*Cre*^*F2rl1*^*+/+*^ PTX mice show a clear loss in nerve fibers in the epidermis that is not seen in the *Pirt*^*Cre*^*F2rl1*^*flox*^ PTX group or vehicle groups. **(B)** To determine the number of intraepidermal nerve fibers (IENF), the number of free nerve endings that cross the dermo-epidermal junction was counted and the IENF density was calculated as the number of fibers per millimeter of epidermal length. The *Pirt*^*Cre*^*F2rl1*^*+/+*^ PTX group showed significantly lower IENF/mm than the *Pirt*^*Cre*^*F2rl1*^*flox*^ PTX group and the *Pirt*^*Cre*^*F2rl1*^*+/+*^ Veh group. **p<0.01, ***p<0.001. For IENF/mm comparisons, Brown-Forsythe and Welch ANOVA with Dunnett’s T3 multiple comparisons test was used. All group mean comparisons were made to the *Pirt*^*Cre*^*F2rl1*^*+/+*^ PTX group.

### Satellite cell gliosis caused by PTX treatment is reduced in mice lacking PAR2 expressed on sensory neurons

Another hallmark of CIPN is satellite glial cell activation in the DRG ^94^. Satellite glial cells surround the cell bodies of DRG neurons and play an important role in intercellular communication with the neurons they surround ^29^. In pathological conditions such as nerve injury or inflammation, satellite cell gliosis is observed as increased expression of glial fibrillary acidic protein (GFAP) ^69, 84, 95^. It is possible that similar to nerve axotomy, the loss in epidermal innervation could lead to satellite cell gliosis ^94^. To determine whether PAR2 expression on sensory neurons is important for satellite cell gliosis as a result of PTX treatment, we dissected the DRG from the mice used in the *in vivo* behavioral assays after completion of experiments. The DRG samples were stained for GFAP to visualize the satellite glial cells and the percentage of neurons surrounded by GFAP-positive satellite glial cells was determined.

In the vehicle-treated animals, few DRG neurons were surrounded by GFAP-positive satellite glial cells. PTX treatment caused an increase in GFAP-positive satellite glial cells in the *Pirt*^*Cre*^*F2rl1*^*+/+*^ mice. However, this increase was not seen in the PTX-treated *Pirt*^*Cre*^*F2rl1*^*flox*^ mice, indicating that PAR2 plays a role in satellite cell gliosis after PTX treatment **(Figure 4A & 4B)**.

**Figure 4:**
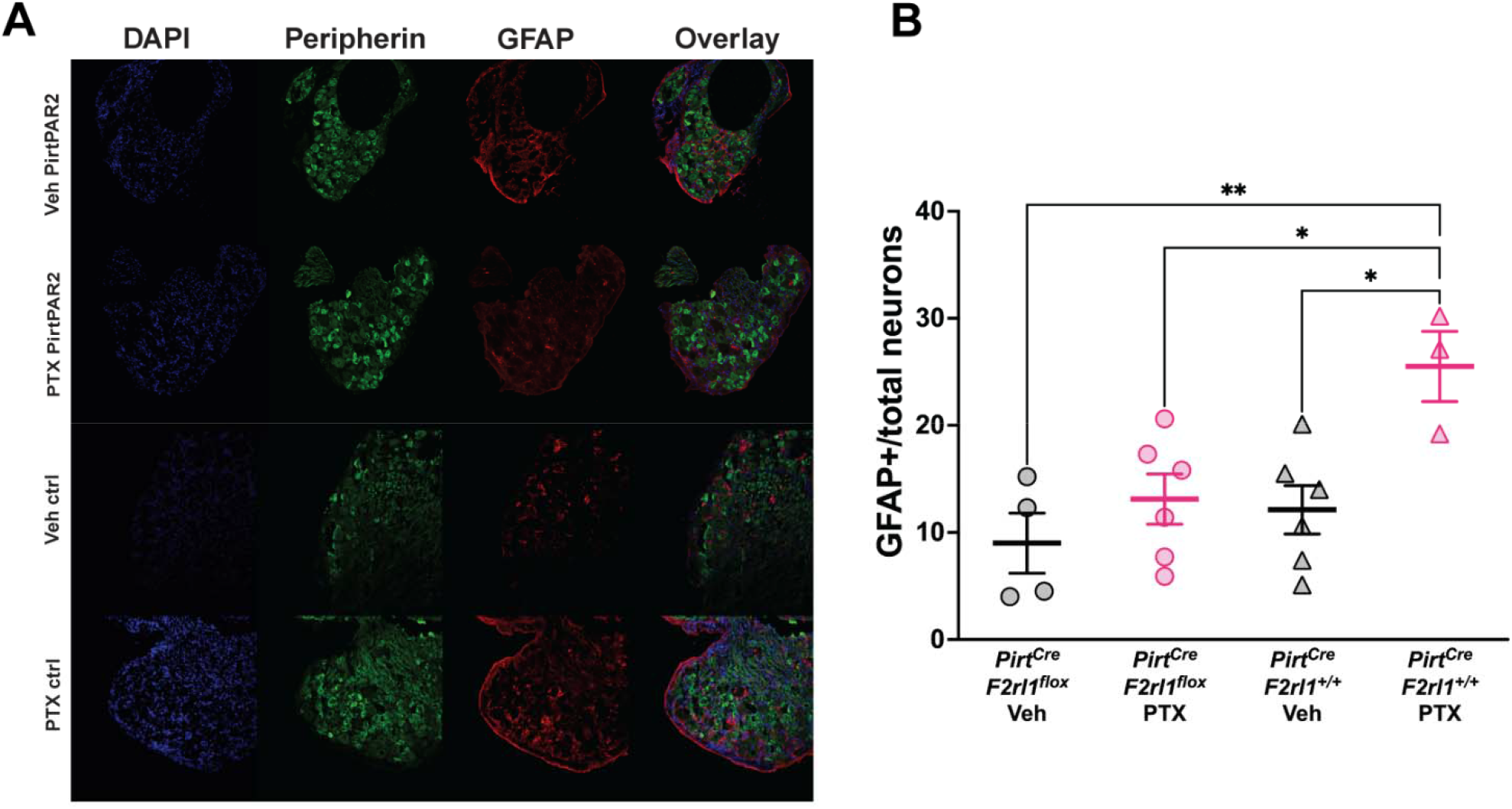
*Pirt*^*Cre*^*F2rl1*^*flox*^ mice are protected from satellite gliosis as a result of paclitaxel treatment. **(A)** Dorsal root ganglion (DRG) was dissected, cryosectioned at 20 µm, and stained for Peripherin (green) to visualize peripheral neurons, GFAP (red) to visualize satellite glial cells, and DAPI (blue) to visualize nuclei. The *Pirt*^*Cre*^*F2rl1*^*+/+*^ PTX mice show clear satellite gliosis as represented by an increase in GFAP expression in the DRG that is not seen in the *Pirt*^*Cre*^*F2rl1*^*flox*^ PTX group or vehicle groups. **(B)** To determine the percentage of neurons that are surrounded by satellite glial cells, the number of neurons that were surrounded by 50% or more by GFAP-positive satellite glial cells was divided by the total number of neurons in the field of view. The *Pirt*^*Cre*^*F2rl1*^*+/+*^ PTX group showed significantly higher of neurons surrounded by GFAP-positive satellite glial cells than all other groups. *p <0.05, **p<0.01. For GFAP^+^/total neurons comparisons, two-way ANOVA with Dunnett’s multiple comparisons test was used. All group mean comparisons were made to the *Pirt*^*Cre*^*F2rl1*^*+/+*^ PTX group.

### A PAR2 antagonist transiently reverses established mechanical hypersensitivity caused by PTX treatment

To determine whether established mechanical hypersensitivity caused by PTX treatment could be reversed by PAR2 signaling inhibition we used a β-arrestin signaling biased PAR2 antagonist that we developed ^78^. C781 was given by intraperitoneal injection and mechanical threshold and facial grimacing was assessed following treatment. Time points were chosen based on previously published pharmacokinetic studies done in mice administered with C781 ^48^. C781 treatment created a transient reversal of mechanical hypersensitivity but did not have an influence on facial grimacing **(Figure 5A & 5B)**.

**Figure 5:**
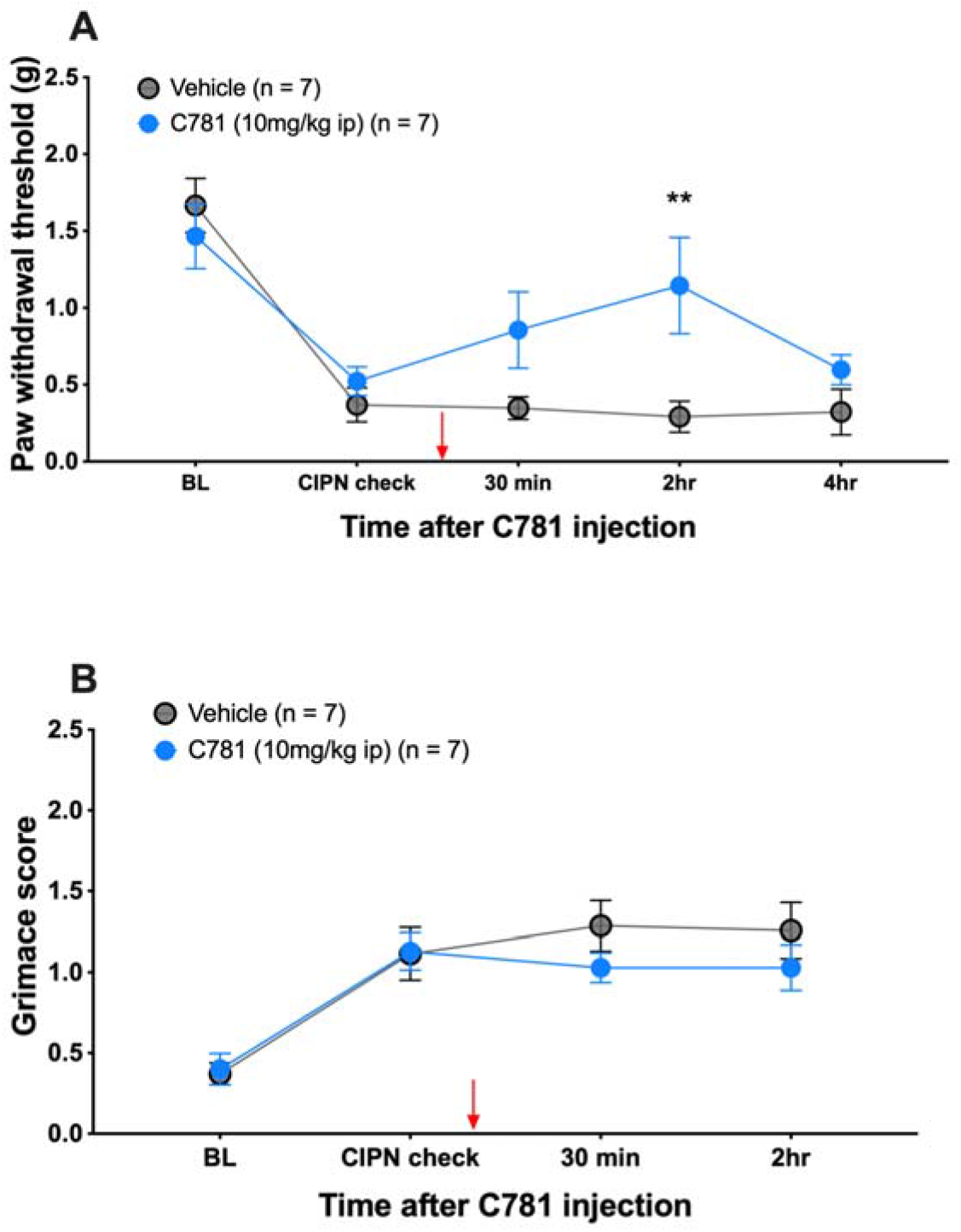
C781 transiently reverses mechanical hypersensitivity in response to paclitaxel treatment. Baseline (BL) paw withdrawal thresholds for male and female ICR mice were obtained before paclitaxel treatment (4 mg/kg IP every other day for 4 days). Before C781 treatment, establishment of CIPN was checked to ensure that animals were mechanically hypersensitive and grimacing. C781 was injected at 10 mg/kg into the intraperitoneal space. **(A)** Mice treated with C781 transiently showed higher mechanical thresholds. **(B)** C781 treatment had no effect on facial grimacing. For the Veh group, n = 3 females and n = 4 males. For the C781 group, n = 4 females and n = 3 males. **p<0.01. For both paw withdrawal threshold and grimace scores, repeated measures two-way ANOVA and Sidak’s multiple comparisons test was used.

## DISCUSSION

In this study we demonstrate that PAR2 expressed on sensory neurons plays an important role in mediating paclitaxel-induced CIPN nociceptive behaviors as well as pathophysiological changes in the skin and DRG that are consistent with neuropathy that is seen in patients. Our findings show that both mechanical hypersensitivity and facial grimacing as a result of paclitaxel treatment are reduced in both global PAR2 knockout mice and mice with PAR2 ablated selectively in sensory neurons. We also demonstrate that paclitaxel-induced skin innervation loss and satellite cell gliosis in the DRG are prevented in *Pirt*^*Cre*^*F2rl1*^*flox*^ mice. Finally, treatment with a PAR2 antagonist, C781, transiently reversed paclitaxel-evoked mechanical hypersensitivity. We conclude that PAR2 expressed in sensory neurons serves as an important mediator of paclitaxel-evoked pain and neuropathy and may be a possible therapeutic target for the treatment of CIPN more broadly.

Previous studies have made a case for PAR2 as an important contributor to CIPN using PAR2 antagonists and global knockout mice, but the cells that express PAR2 to mediate these effects have been unclear ^15^. A key contribution of our work is clarifying that PAR2 expressed by sensory neurons is the site of action for the role of PAR2 in CIPN, at least for paclitaxel treatment. Previous studies using sensory neuron ablation of PAR2 have shown that sensory neuronal PAR2 is critical for pain caused by injection of exogenous proteases, pain caused by inflammatory mediators, pain caused by proteases released from immune and cancer cells and certain types of visceral pain ^31^. In the DRG of mice, PAR2 is expressed by a specific subset of neurons that express markers that are consistent with these neurons innervating the outermost layer of the skin as free nerve endings in the epidermis ^31, 51, 63, 66^. Our finding that intraepidermal nerve fiber loss is abrogated with sensory neuron-specific ablation of PAR2 is consistent with this previous literature. Although PAR2 is only expressed by a small subset of sensory neurons, we noted a nearly complete loss of satellite cell gliosis caused by paclitaxel treatment in the DRGs of mice with sensory neuron cKO of PAR2. The reasons for this effect are currently unclear, but suggest that PAR2 is a key mediator of cascading signaling events that lead to a broad satellite cell gliosis that eventually envelops neurons that likely do not express PAR2. This could be a mechanism through which a neuropathy that usually first affects long axons that innervate the epidermis can spread to other neuronal subtypes causing spontaneous pain in CIPN patients.

Paclitaxel is considered one of the most neurotoxic chemotherapeutics, causing one of the highest CIPN rates in patients ^67^. This is in line with preclinical CIPN models, in which paclitaxel or cisplatin treatment in mice showed the highest efficacy in causing CIPN ^25^. Although many clinical trials have investigated potential therapies for the prevention or treatment of CIPN, as of 2020, only duloxetine has emerged as a moderately recommended treatment by the American Society of Clinical Oncology (ASCO) clinical practice guidelines ^20, 35, 59, 98^. However, duloxetine has been shown to be more effective in alleviating CIPN as a result of platinum-based therapies like cisplatin rather than taxane-treated patients, suggesting a difference in CIPN mechanism ^81^. Most clinical trial studies have been based on evidence from other types of neuropathies (e.g., diabetic neuropathy) but the mechanism(s) through which CIPN develops differentially from other chronic neuropathic pain conditions, and also how these mechanisms may differ between various chemotherapeutic agents, remain unclear ^10, 38^. CIPN can develop through multiple targets, and while the neurotoxic mechanisms that initiate and maintain CIPN potentially overlap with its cytotoxic mechanisms, it is imperative that CIPN-blocking drugs do not function at the expense of the antitumor efficacy of chemotherapeutics ^10, 38^. PAR2 has been associated with cancer where it is thought to play a role in promoting oncogenesis and cancer cell invasiveness ^83, 92^. While PAR2 antagonists as a treatment for CIPN would require testing on specific cancer types, we would not expect that blocking PAR2 would cause interference with the efficacy of chemotherapeutics given the existing literature on PAR2 and cancer ^41^.

Paclitaxel has been shown to cause immune cell infiltration and activation in the skin and in the DRG, including increased infiltration and activation of macrophages, mast cells, and neutrophils ^15, 26, 58, 71, 72, 100^. PAR2 thus emerged as a possible mediator of CIPN as these activated immune cells release proteases which act on PAR2, causing sensitization of sensory neurons and hyperalgesia ^7, 15, 68, 75, 86^. Macrophages are known to release PAR2-activating proteases including elastase and cathepsin S, and the use of minocycline, a macrophage inhibitor, prevents macrophage recruitment into the DRG as well as mechanical hypersensitivity caused by paclitaxel treatment ^9, 11, 12, 18, 57^. Similar results were also seen with quercetin, a mast cell stabilizer ^26^.

Our work here fits with previous studies on signaling within sensory neurons that causes many features of CIPN, including mitochondrial and lysosomal dysfunction and altered translation regulation that causes nociceptor hyperexcitability ^19, 22^. The model emerging from the present experiments, and previous work on mechanisms of paclitaxel-evoked pain, is that PAR2 activation on nociceptors induces β-arrestin-mediated mitogen activated protein kinase (MAPK) activation including phosphorylation of extracellular regulated protein kinase (ERK) and mitogen activated protein kinase interacting kinase (MNK) ^87^. This then induces changes in gene expression driven by phosphorylation of proteins that control translation of mRNAs that are involved in nociceptor hyperexcitability, and lysosomal and mitochondrial dysfunction via increased translation of RagA and altered mechanistic target of rapamycin (mTOR) signaling at the surface of lysosomes ^62^. This pathology likely starts in neurons that innervate the skin, in particular the skin of the hands and feet, but it may spread to a broader subset of neurons as paclitaxel causes hyperexcitability in many nociceptor populations both in mice and in humans (**Figure 6**). It will be important to understand how this broader effect of paclitaxel spreads and what role PAR2 plays in initiating this cascade of signaling events. As mentioned above, one testable hypothesis is satellite cell gliosis which is well known to occur in CIPN mouse models. Interestingly we recently showed that meteorin, which is thought to target satellite glial cells, can both prevent and reverse paclitaxel-evoked pain and signs of neuropathy ^77^.

**Figure 6:**
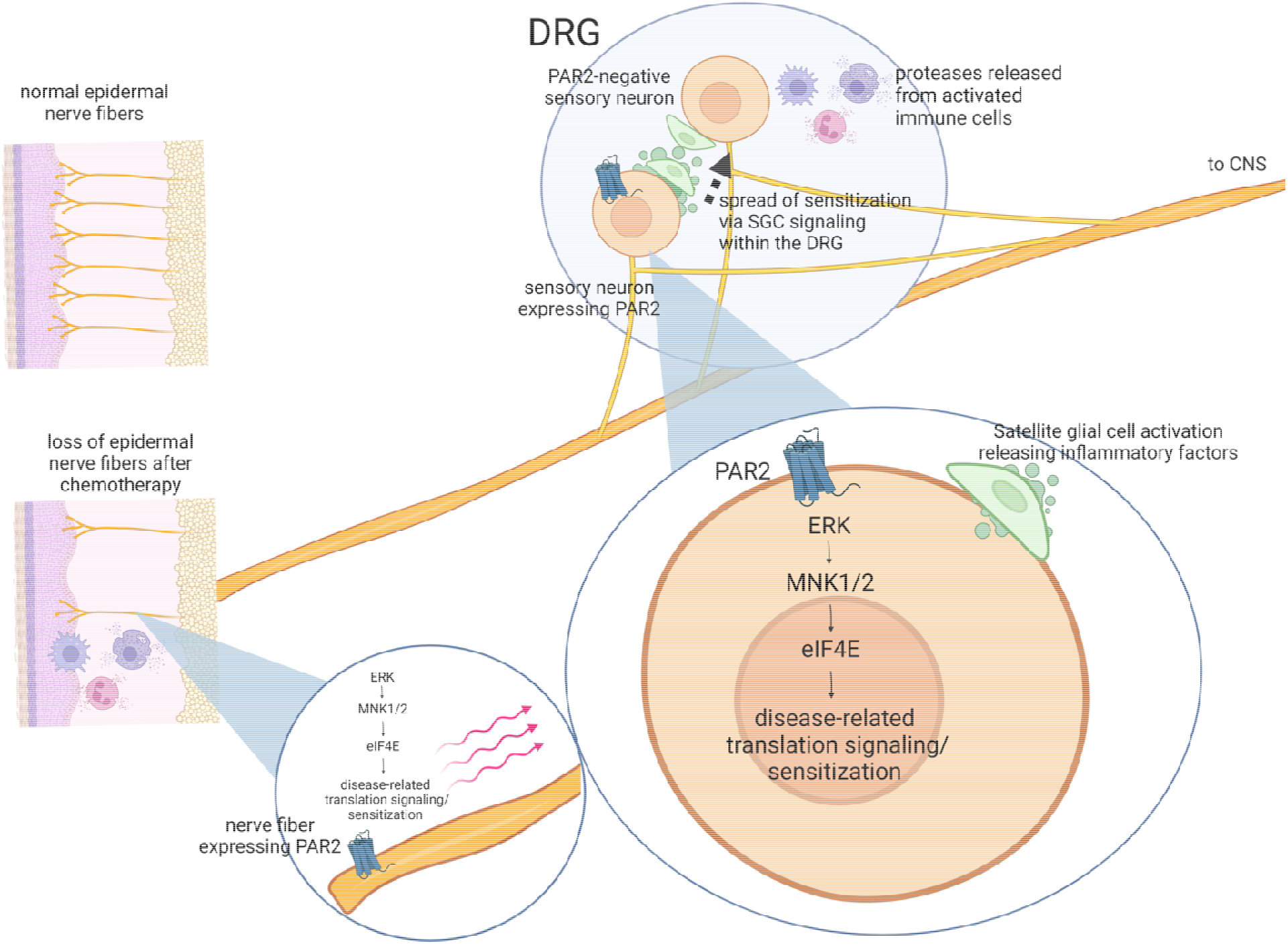
Summary diagram of potential role of PAR2 and downstream signaling in paclitaxel-induced neuropathy. The figure summarizes how PAR2 signaling on a specific subset of nociceptors might lead to loss of epidermal nerve fibers, satellite cell gliosis and pain. A key feature of the model is that paclitaxel-induced changes may begin with a small subset of sensory fibers, but gene expression changes in DRG cell bodies driven by PAR2 activation could cause spread of satellite cell gliosis causing sensitization of additional subsets of DRG nociceptors.

C781 only transiently reversed the mechanical hypersensitivity caused by paclitaxel in mice and had no effect on grimacing. This transient effect is likely attributable to the pharmacokinetics of C781. We previously published that C781 achieves a rapid spike in plasma concentration with systemic administration and then is likely excreted via the kidneys, although we cannot rule out excretion via feces ^48^. It is unlikely that the compound is metabolized in the liver because C781 has a long half-life in liver microsomes. Our pharmacological study should be viewed as proof of concept for the efficacy of a PAR2 antagonist that selectively blocks β-arrestin signaling downstream of the receptor for the treatment of CIPN ^78^. Future development of similar PAR2 antagonists with a more favorable pharmacokinetic profile will allow for longer target coverage *in vivo*. Based on our findings with PAR2 cKO mice, we hypothesize that such a molecule would have efficacy against pain endpoints, as well as reversing signs of small fiber neuropathy such as loss of intraepidermal nerve fibers.

In conclusion, our work provides clarity on how PAR2 is involved in CIPN. Sensory neuron expressed PAR2 is responsible for mechanical hypersensitivity, facial grimacing, intraepidermal nerve fiber loss and satellite cell gliosis caused by paclitaxel in mice. Our data with C781 suggests, in agreement with previous studies, that PAR2 can be targeted pharmacologically for the alleviation of CIPN pain even after chemotherapeutic treatment.

## Acknowledgements

This work was supported by NIH grants NS098826, NS065926, and AI140257.

## References Cited

1. Abal M, Andreu JM, Barasoain I: Taxanes: microtubule and centrosome targets, and cell cycle dependent mechanisms of action. Curr Cancer Drug Targets 3:193–203, 2003

2. Adams MN, Ramachandran R, Yau MK, Suen JY, Fairlie DP, Hollenberg MD, Hooper JD: Structure, function and pathophysiology of protease activated receptors. Pharmacol Ther 130:248–82, 2011

3. Agalave NM, Mody PH, Szabo-Pardi TA, Jeong HS, Burton MD: Neuroimmune Consequences of eIF4E Phosphorylation on Chemotherapy-Induced Peripheral Neuropathy. Front Immunol 12:642420, 2021

4. Akintola T, Raver C, Studlack P, Uddin O, Masri R, Keller A: The grimace scale reliably assesses chronic pain in a rodent model of trigeminal neuropathic pain. Neurobiol Pain 2:13–7, 2017

5. Amos LA, Lowe J: How Taxol stabilises microtubule structure. Chem Biol 6:R65–9, 1999

6. Banach M, Juranek JK, Zygulska AL: Chemotherapy-induced neuropathies-a growing problem for patients and health care providers. Brain Behav 7:e00558, 2017

7. Bao Y, Hou W, Hua B: Protease-activated receptor 2 signalling pathways: a role in pain processing. Expert Opin Ther Targets 18:15–27, 2014

8. Bhatnagar B, Gilmore S, Goloubeva O, Pelser C, Medeiros M, Chumsri S, Tkaczuk K, Edelman M, Bao T: Chemotherapy dose reduction due to chemotherapy induced peripheral neuropathy in breast cancer patients receiving chemotherapy in the neoadjuvant or adjuvant settings: a single-center experience. Springerplus 3:366, 2014

9. Boyette-Davis J, Xin W, Zhang H, Dougherty PM: Intraepidermal nerve fiber loss corresponds to the development of taxol-induced hyperalgesia and can be prevented by treatment with minocycline. Pain 152:308–13, 2011

10. Carozzi VA, Canta A, Chiorazzi A: Chemotherapy-induced peripheral neuropathy: What do we know about mechanisms? Neurosci Lett 596:90–107, 2015

11. Cata JP, Weng HR, Dougherty PM: The effects of thalidomide and minocycline on taxol-induced hyperalgesia in rats. Brain Res 1229:100–10, 2008

12. Cattaruzza F, Lyo V, Jones E, Pham D, Hawkins J, Kirkwood K, Valdez-Morales E, Ibeakanma C, Vanner SJ, Bogyo M, Bunnett NW: Cathepsin S is activated during colitis and causes visceral hyperalgesia by a PAR2-dependent mechanism in mice. Gastroenterology 141:1864-74 e1-3, 2011

13. Cavaletti G, Cavalletti E, Oggioni N, Sottani C, Minoia C, D’Incalci M, Zucchetti M, Marmiroli P, Tredici G: Distribution of paclitaxel within the nervous system of the rat after repeated intravenous administration. Neurotoxicology 21:389–93, 2000

14. Chaplan SR, Bach FW, Pogrel JW, Chung JM, Yaksh TL: Quantitative assessment of tactile allodynia in the rat paw. J Neurosci Methods 53:55–63, 1994

15. Chen Y, Yang C, Wang ZJ: Proteinase-activated receptor 2 sensitizes transient receptor potential vanilloid 1, transient receptor potential vanilloid 4, and transient receptor potential ankyrin 1 in paclitaxel-induced neuropathic pain. Neuroscience 193:440–51, 2011

16. Chine VB, Au NPB, Kumar G, Ma CHE: Targeting Axon Integrity to Prevent Chemotherapy-Induced Peripheral Neuropathy. Mol Neurobiol 56:3244–59, 2019

17. de Moor JS, Mariotto AB, Parry C, Alfano CM, Padgett L, Kent EE, Forsythe L, Scoppa S, Hachey M, Rowland JH: Cancer survivors in the United States: prevalence across the survivorship trajectory and implications for care. Cancer Epidemiol Biomarkers Prev 22:561–70, 2013

18. Dollery CM, Owen CA, Sukhova GK, Krettek A, Shapiro SD, Libby P: Neutrophil elastase in human atherosclerotic plaques: production by macrophages. Circulation 107:2829–36, 2003

19. Duggett NA, Griffiths LA, McKenna OE, de Santis V, Yongsanguanchai N, Mokori EB, Flatters SJ: Oxidative stress in the development, maintenance and resolution of paclitaxel-induced painful neuropathy. Neuroscience 333:13–26, 2016

20. Farshchian N, Alavi A, Heydarheydari S, Moradian N: Comparative study of the effects of venlafaxine and duloxetine on chemotherapy-induced peripheral neuropathy. Cancer Chemother Pharmacol 82:787–93, 2018

21. Fellner S, Bauer B, Miller DS, Schaffrik M, Fankhanel M, Spruss T, Bernhardt G, Graeff C, Farber L, Gschaidmeier H, Buschauer A, Fricker G: Transport of paclitaxel (Taxol) across the blood-brain barrier in vitro and in vivo. J Clin Invest 110:1309–18, 2002

22. Flatters SJL, Bennett GJ: Studies of peripheral sensory nerves in paclitaxel-induced painful peripheral neuropathy: evidence for mitochondrial dysfunction. Pain 122:245–57, 2006

23. Flatters SJL, Dougherty PM, Colvin LA: Clinical and preclinical perspectives on Chemotherapy-Induced Peripheral Neuropathy (CIPN): a narrative review. Br J Anaesth 119:737–49, 2017

24. Fradkin M, Batash R, Elmaleh S, Debi R, Schaffer P, Schaffer M, Asna N: Management of Peripheral Neuropathy Induced by Chemotherapy. Curr Med Chem 26:4698–708, 2019

25. Gadgil S, Ergun M, van den Heuvel SA, van der Wal SE, Scheffer GJ, Hooijmans CR: A systematic summary and comparison of animal models for chemotherapy induced (peripheral) neuropathy (CIPN). PLoS One 14:e0221787, 2019

26. Gao W, Zan Y, Wang ZJ, Hu XY, Huang F: Quercetin ameliorates paclitaxel-induced neuropathic pain by stabilizing mast cells, and subsequently blocking PKCepsilon-dependent activation of TRPV1. Acta Pharmacol Sin 37:1166–77, 2016

27. Glare PA, Davies PS, Finlay E, Gulati A, Lemanne D, Moryl N, Oeffinger KC, Paice JA, Stubblefield MD, Syrjala KL: Pain in cancer survivors. J Clin Oncol 32:1739–47, 2014

28. Grisold W, Cavaletti G, Windebank AJ: Peripheral neuropathies from chemotherapeutics and targeted agents: diagnosis, treatment, and prevention. Neuro Oncol 14 Suppl 4:iv45–54, 2012

29. Hanani M: Intercellular communication in sensory ganglia by purinergic receptors and gap junctions: implications for chronic pain. Brain Res 1487:183–91, 2012

30. Hassler SN, Ahmad FB, Burgos-Vega CC, Boitano S, Vagner J, Price TJ, Dussor G: Protease activated receptor 2 (PAR2) activation causes migraine-like pain behaviors in mice. Cephalalgia 39:111–22, 2019

31. Hassler SN, Kume M, Mwirigi J, Ahmad A, Shiers S, Wangzhou A, Ray P, Belugin SN, Naik DK, Burton MD, Vagner J, Boitano S, Akopian AN, Dussor G, Price TJ: The cellular basis of protease activated receptor type 2 (PAR2) evoked mechanical and affective pain. JCI Insight, 2020

32. Hassler SN, Kume M, Mwirigi JM, Ahmad A, Shiers S, Wangzhou A, Ray PR, Belugin SN, Naik DK, Burton MD, Vagner J, Boitano S, Akopian AN, Dussor G, Price TJ: The cellular basis of protease-activated receptor 2-evoked mechanical and affective pain. JCI Insight 5, 2020

33. Hershman DL, Lacchetti C, Dworkin RH, Lavoie Smith EM, Bleeker J, Cavaletti G, Chauhan C, Gavin P, Lavino A, Lustberg MB, Paice J, Schneider B, Smith ML, Smith T, Terstriep S, Wagner-Johnston N, Bak K, Loprinzi CL, American Society of Clinical O: Prevention and management of chemotherapy-induced peripheral neuropathy in survivors of adult cancers: American Society of Clinical Oncology clinical practice guideline. J Clin Oncol 32:1941–67, 2014

34. Heuberger DM, Schuepbach RA: Protease-activated receptors (PARs): mechanisms of action and potential therapeutic modulators in PAR-driven inflammatory diseases. Thromb J 17:4, 2019

35. Hirayama Y, Ishitani K, Sato Y, Iyama S, Takada K, Murase K, Kuroda H, Nagamachi Y, Konuma Y, Fujimi A, Sagawa T, Ono K, Horiguchi H, Terui T, Koike K, Kusakabe T, Sato T, Takimoto R, Kobune M, Kato J: Effect of duloxetine in Japanese patients with chemotherapy-induced peripheral neuropathy: a pilot randomized trial. Int J Clin Oncol 20:866–71, 2015

36. Hopkins HL, Duggett NA, Flatters SJL: Chemotherapy-induced painful neuropathy: pain-like behaviours in rodent models and their response to commonly used analgesics. Curr Opin Support Palliat Care 10:119–28, 2016

37. Hornick JE, Bader JR, Tribble EK, Trimble K, Breunig JS, Halpin ES, Vaughan KT, Hinchcliffe EH: Live-cell analysis of mitotic spindle formation in taxol-treated cells. Cell Motil Cytoskeleton 65:595–613, 2008

38. Hu S, Huang KM, Adams EJ, Loprinzi CL, Lustberg MB: Recent Developments of Novel Pharmacologic Therapeutics for Prevention of Chemotherapy-Induced Peripheral Neuropathy. Clin Cancer Res 25:6295–301, 2019

39. Inyang KE, McDougal TA, Ramirez ED, Williams M, Laumet G, Kavelaars A, Heijnen CJ, Burton M, Dussor G, Price TJ: Alleviation of paclitaxel-induced mechanical hypersensitivity and hyperalgesic priming with AMPK activators in male and female mice. Neurobiol Pain 6:100037, 2019

40. Jemal A, Center MM, DeSantis C, Ward EM: Global patterns of cancer incidence and mortality rates and trends. Cancer Epidemiol Biomarkers Prev 19:1893–907, 2010

41. Jiang Y, Yau MK, Lim J, Wu KC, Xu W, Suen JY, Fairlie DP: A Potent Antagonist of Protease-Activated Receptor 2 That Inhibits Multiple Signaling Functions in Human Cancer Cells. J Pharmacol Exp Ther 364:246–57, 2018

42. Jimenez-Vargas NN, Pattison LA, Zhao P, Lieu T, Latorre R, Jensen DD, Castro J, Aurelio L, Le GT, Flynn B, Herenbrink CK, Yeatman HR, Edgington-Mitchell L, Porter CJH, Halls ML, Canals M, Veldhuis NA, Poole DP, McLean P, Hicks GA, Scheff N, Chen E, Bhattacharya A, Schmidt BL, Brierley SM, Vanner SJ, Bunnett NW: Protease-activated receptor-2 in endosomes signals persistent pain of irritable bowel syndrome. Proc Natl Acad Sci U S A 115:E7438–E47, 2018

43. Kim AY, Tang Z, Liu Q, Patel KN, Maag D, Geng Y, Dong X: Pirt, a phosphoinositide-binding protein, functions as a regulatory subunit of TRPV1. Cell 133:475–85, 2008

44. Kim YS, Anderson M, Park K, Zheng Q, Agarwal A, Gong C, Saijilafu, Young L, He S, LaVinka PC, Zhou F, Bergles D, Hanani M, Guan Y, Spray DC, Dong X: Coupled Activation of Primary Sensory Neurons Contributes to Chronic Pain. Neuron 91:1085–96, 2016

45. Ko MH, Hu ME, Hsieh YL, Lan CT, Tseng TJ: Peptidergic intraepidermal nerve fibers in the skin contribute to the neuropathic pain in paclitaxel-induced peripheral neuropathy. Neuropeptides 48:109–17, 2014

46. Kopruszinski CM, Thornton P, Arnold J, Newton P, Lowne D, Navratilova E, Swiokla J, Dodick DW, Dobson C, Gurrell I, Chessell IP, Porreca F: Characterization and preclinical evaluation of a protease activated receptor 2 (PAR2) monoclonal antibody as a preventive therapy for migraine. Cephalalgia 40:1535–50, 2020

47. Kumar P, Lau CS, Mathur M, Wang P, DeFea KA: Differential effects of beta-arrestins on the internalization, desensitization and ERK1/2 activation downstream of protease activated receptor-2. Am J Physiol Cell Physiol 293:C346–57, 2007

48. Kume M, Ahmad A, Shiers S, Burton MD, DeFea KA, Vagner J, Dussor G, Boitano S, Price TJ: C781, a beta-arrestin biased antagonist at protease-activated receptor-2 (PAR2), displays in vivo efficacy against protease-induced pain in mice. J Pain, 2022

49. Lam DK, Dang D, Zhang J, Dolan JC, Schmidt BL: Novel animal models of acute and chronic cancer pain: a pivotal role for PAR2. J Neurosci 32:14178–83, 2012

50. Lam DK, Dang D, Flynn AN, Hardt M, Schmidt BL: TMPRSS2, a novel membrane-anchored mediator in cancer pain. Pain 156:923–30, 2015

51. LaMotte RH, Dong X, Ringkamp M: Sensory neurons and circuits mediating itch. Nat Rev Neurosci 15:19–31, 2014

52. Langford DJ, Bailey AL, Chanda ML, Clarke SE, Drummond TE, Echols S, Glick S, Ingrao J, Klassen-Ross T, Lacroix-Fralish ML, Matsumiya L, Sorge RE, Sotocinal SG, Tabaka JM, Wong D, van den Maagdenberg AM, Ferrari MD, Craig KD, Mogil JS: Coding of facial expressions of pain in the laboratory mouse. Nature methods 7:447–9, 2010

53. Lauria G, Cornblath DR, Johansson O, McArthur JC, Mellgren SI, Nolano M, Rosenberg N, Sommer C, European Federation of Neurological S: EFNS guidelines on the use of skin biopsy in the diagnosis of peripheral neuropathy. Eur J Neurol 12:747–58, 2005

54. Leach MC, Klaus K, Miller AL, Scotto di Perrotolo M, Sotocinal SG, Flecknell PA: The assessment of post-vasectomy pain in mice using behaviour and the Mouse Grimace Scale. PLoS One 7:e35656, 2012

55. Li CL, Li KC, Wu D, Chen Y, Luo H, Zhao JR, Wang SS, Sun MM, Lu YJ, Zhong YQ, Hu XY, Hou R, Zhou BB, Bao L, Xiao HS, Zhang X: Somatosensory neuron types identified by high-coverage single-cell RNA-sequencing and functional heterogeneity. Cell research 26:83–102, 2016

56. Lipton RB, Apfel SC, Dutcher JP, Rosenberg R, Kaplan J, Berger A, Einzig AI, Wiernik P, Schaumburg HH: Taxol produces a predominantly sensory neuropathy. Neurology 39:368–73, 1989

57. Liu CC, Lu N, Cui Y, Yang T, Zhao ZQ, Xin WJ, Liu XG: Prevention of paclitaxel-induced allodynia by minocycline: Effect on loss of peripheral nerve fibers and infiltration of macrophages in rats. Mol Pain 6:76, 2010

58. Liu XJ, Zhang Y, Liu T, Xu ZZ, Park CK, Berta T, Jiang D, Ji RR: Nociceptive neurons regulate innate and adaptive immunity and neuropathic pain through MyD88 adapter. Cell Res 24:1374–7, 2014

59. Loprinzi CL, Lacchetti C, Bleeker J, Cavaletti G, Chauhan C, Hertz DL, Kelley MR, Lavino A, Lustberg MB, Paice JA, Schneider BP, Lavoie Smith EM, Smith ML, Smith TJ, Wagner-Johnston N, Hershman DL: Prevention and Management of Chemotherapy-Induced Peripheral Neuropathy in Survivors of Adult Cancers: ASCO Guideline Update. J Clin Oncol 38:3325–48, 2020

60. Matsumiya LC, Sorge RE, Sotocinal SG, Tabaka JM, Wieskopf JS, Zaloum A, King OD, Mogil JS: Using the Mouse Grimace Scale to reevaluate the efficacy of postoperative analgesics in laboratory mice. J Am Assoc Lab Anim Sci 51:42–9, 2012

61. McCulloch K, McGrath S, Huesa C, Dunning L, Litherland G, Crilly A, Hultin L, Ferrell WR, Lockhart JC, Goodyear CS: Rheumatic Disease: Protease-Activated Receptor-2 in Synovial Joint Pathobiology. Front Endocrinol (Lausanne) 9:257, 2018

62. Megat S, Ray PR, Moy JK, Lou TF, Barragan-Iglesias P, Li Y, Pradhan G, Wanghzou A, Ahmad A, Burton MD, North RY, Dougherty PM, Khoutorsky A, Sonenberg N, Webster KR, Dussor G, Campbell ZT, Price TJ: Nociceptor Translational Profiling Reveals the Ragulator-Rag GTPase Complex as a Critical Generator of Neuropathic Pain. J Neurosci 39:393–411, 2019

63. Meixiong J, Dong X: Mas-Related G Protein-Coupled Receptors and the Biology of Itch Sensation. Annu Rev Genet 51:103–21, 2017

64. Miller KD, Nogueira L, Mariotto AB, Rowland JH, Yabroff KR, Alfano CM, Jemal A, Kramer JL, Siegel RL: Cancer treatment and survivorship statistics, 2019. CA Cancer J Clin 69:363–85, 2019

65. Miltenburg NC, Boogerd W: Chemotherapy-induced neuropathy: A comprehensive survey. Cancer Treat Rev 40:872–82, 2014

66. Mishra SK, Hoon MA: The cells and circuitry for itch responses in mice. Science 340:968–71, 2013

67. Molassiotis A, Cheng HL, Lopez V, Au JSK, Chan A, Bandla A, Leung KT, Li YC, Wong KH, Suen LKP, Chan CW, Yorke J, Farrell C, Sundar R: Are we mis-estimating chemotherapy-induced peripheral neuropathy? Analysis of assessment methodologies from a prospective, multinational, longitudinal cohort study of patients receiving neurotoxic chemotherapy. BMC Cancer 19:132, 2019

68. Mrozkova P, Palecek J, Spicarova D: The role of protease-activated receptor type 2 in nociceptive signaling and pain. Physiol Res 65:357–67, 2016

69. Nascimento DS, Castro-Lopes JM, Moreira Neto FL: Satellite glial cells surrounding primary afferent neurons are activated and proliferate during monoarthritis in rats: is there a role for ATF3? PLoS One 9:e108152, 2014

70. Park SB, Goldstein D, Krishnan AV, Lin CS, Friedlander ML, Cassidy J, Koltzenburg M, Kiernan MC: Chemotherapy-induced peripheral neurotoxicity: a critical analysis. CA Cancer J Clin 63:419–37, 2013

71. Peters CM, Jimenez-Andrade JM, Jonas BM, Sevcik MA, Koewler NJ, Ghilardi JR, Wong GY, Mantyh PW: Intravenous paclitaxel administration in the rat induces a peripheral sensory neuropathy characterized by macrophage infiltration and injury to sensory neurons and their supporting cells. Exp Neurol 203:42–54, 2007

72. Peters CM, Jimenez-Andrade JM, Kuskowski MA, Ghilardi JR, Mantyh PW: An evolving cellular pathology occurs in dorsal root ganglia, peripheral nerve and spinal cord following intravenous administration of paclitaxel in the rat. Brain Res 1168:46–59, 2007

73. Ramachandran R, Hollenberg MD: Proteinases and signalling: pathophysiological and therapeutic implications via PARs and more. Br J Pharmacol 153 Suppl 1:S263–82, 2008

74. Ramachandran R, Noorbakhsh F, Defea K, Hollenberg MD: Targeting proteinase-activated receptors: therapeutic potential and challenges. Nat Rev Drug Discov 11:69–86, 2012

75. Sakamoto A, Andoh T, Kuraishi Y: Involvement of mast cells and proteinase-activated receptor 2 in oxaliplatin-induced mechanical allodynia in mice. Pharmacol Res 105:84–92, 2016

76. Salat K: Chemotherapy-induced peripheral neuropathy: part 1-current state of knowledge and perspectives for pharmacotherapy. Pharmacol Rep 72:486–507, 2020

77. Sankaranarayanan I, Tavares-Ferreira D, He L, Kume M, Mwirigi JM, Madsen TM, Petersen KA, Munro G, Price TJ: Meteorin Alleviates Paclitaxel-Induced Peripheral Neuropathic Pain in Mice. J Pain, 2022

78. Schiff HV, Rivas CM, Pederson WP, Sandoval E, Gillman S, Prisco J, Kume M, Dussor G, Vagner J, Ledford JG, Price TJ, DeFea KA, Boitano S: beta-arrestin-biased proteinase-activated receptor-2 antagonist C781 limits allergen-induced airway hyperresponsiveness and inflammation. Br J Pharmacol, 2022

79. Seretny M, Currie GL, Sena ES, Ramnarine S, Grant R, MacLeod MR, Colvin LA, Fallon M: Incidence, prevalence, and predictors of chemotherapy-induced peripheral neuropathy: A systematic review and meta-analysis. Pain 155:2461–70, 2014

80. Siau C, Xiao W, Bennett GJ: Paclitaxel- and vincristine-evoked painful peripheral neuropathies: loss of epidermal innervation and activation of Langerhans cells. Exp Neurol 201:507–14, 2006

81. Smith EM, Pang H, Cirrincione C, Fleishman S, Paskett ED, Ahles T, Bressler LR, Fadul CE, Knox C, Le-Lindqwister N, Gilman PB, Shapiro CL, Alliance for Clinical Trials in O: Effect of duloxetine on pain, function, and quality of life among patients with chemotherapy-induced painful peripheral neuropathy: a randomized clinical trial. JAMA 309:1359–67, 2013

82. Speck RM, Sammel MD, Farrar JT, Hennessy S, Mao JJ, Stineman MG, DeMichele A: Impact of chemotherapy-induced peripheral neuropathy on treatment delivery in nonmetastatic breast cancer. J Oncol Pract 9:e234–40, 2013

83. Sun L, Li PB, Yao YF, Xiu AY, Peng Z, Bai YH, Gao YJ: Proteinase-activated receptor 2 promotes tumor cell proliferation and metastasis by inducing epithelial-mesenchymal transition and predicts poor prognosis in hepatocellular carcinoma. World J Gastroenterol 24:1120–33, 2018

84. Takeda M, Tanimoto T, Kadoi J, Nasu M, Takahashi M, Kitagawa J, Matsumoto S: Enhanced excitability of nociceptive trigeminal ganglion neurons by satellite glial cytokine following peripheral inflammation. Pain 129:155–66, 2007

85. Tanay MAL, Armes J, Ream E: The experience of chemotherapy-induced peripheral neuropathy in adult cancer patients: a qualitative thematic synthesis. Eur J Cancer Care (Engl) 26, 2017

86. Tian L, Fan T, Zhou N, Guo H, Zhang W: Role of PAR2 in regulating oxaliplatin-induced neuropathic pain via TRPA1. Transl Neurosci 6:111–6, 2015

87. Tillu DV, Hassler SN, Burgos-Vega CC, Quinn TL, Sorge RE, Dussor G, Boitano S, Vagner J, Price TJ: Protease-activated receptor 2 activation is sufficient to induce the transition to a chronic pain state. Pain 156:859–67, 2015

88. Tofthagen C: Patient perceptions associated with chemotherapy-induced peripheral neuropathy. Clin J Oncol Nurs 14:E22–8, 2010

89. Toma W, Kyte SL, Bagdas D, Alkhlaif Y, Alsharari SD, Lichtman AH, Chen ZJ, Del Fabbro E, Bigbee JW, Gewirtz DA, Damaj MI: Effects of paclitaxel on the development of neuropathy and affective behaviors in the mouse. Neuropharmacology 117:305–15, 2017

90. Tu NH, Jensen DD, Anderson BM, Chen E, Jimenez-Vargas NN, Scheff NN, Inoue K, Tran HD, Dolan JC, Meek TA, Hollenberg MD, Liu CZ, Vanner SJ, Janal MN, Bunnett NW, Edgington-Mitchell LE, Schmidt BL: Legumain Induces Oral Cancer Pain by Biased Agonism of Protease-Activated Receptor-2. J Neurosci 41:193–210, 2021

91. Tuttle AH, Molinaro MJ, Jethwa JF, Sotocinal SG, Prieto JC, Styner MA, Mogil JS, Zylka MJ: A deep neural network to assess spontaneous pain from mouse facial expressions. Mol Pain 14:1744806918763658, 2018

92. Ungefroren H, Witte D, Rauch BH, Settmacher U, Lehnert H, Gieseler F, Kaufmann R: Proteinase-Activated Receptor 2 May Drive Cancer Progression by Facilitating TGF-beta Signaling. Int J Mol Sci 18, 2017

93. Wani MC, Taylor HL, Wall ME, Coggon P, McPhail AT: Plant antitumor agents. VI. The isolation and structure of taxol, a novel antileukemic and antitumor agent from Taxus brevifolia. J Am Chem Soc 93:2325–7, 1971

94. Warwick RA, Hanani M: The contribution of satellite glial cells to chemotherapy-induced neuropathic pain. Eur J Pain 17:571–80, 2013

95. Woodham P, Anderson PN, Nadim W, Turmaine M: Satellite cells surrounding axotomised rat dorsal root ganglion cells increase expression of a GFAP-like protein. Neurosci Lett 98:8–12, 1989

96. World Health O: The selection and use of essential medicines. World Health Organ Tech Rep Ser:i-xiv, 1-219, back cover, 2014

97. Wozniak KM, Vornov JJ, Wu Y, Liu Y, Carozzi VA, Rodriguez-Menendez V, Ballarini E, Alberti P, Pozzi E, Semperboni S, Cook BM, Littlefield BA, Nomoto K, Condon K, Eckley S, DesJardins C, Wilson L, Jordan MA, Feinstein SC, Cavaletti G, Polydefkis M, Slusher BS: Peripheral Neuropathy Induced by Microtubule-Targeted Chemotherapies: Insights into Acute Injury and Long-term Recovery. Cancer Res 78:817–29, 2018

98. Yang YH, Lin JK, Chen WS, Lin TC, Yang SH, Jiang JK, Chang SC, Lan YT, Lin CC, Yen CC, Tzeng CH, Wang WS, Chiang HL, Teng CJ, Teng HW: Duloxetine improves oxaliplatin-induced neuropathy in patients with colorectal cancer: an open-label pilot study. Support Care Cancer 20:1491–7, 2012

99. Zajaczkowska R, Kocot-Kepska M, Leppert W, Wrzosek A, Mika J, Wordliczek J: Mechanisms of Chemotherapy-Induced Peripheral Neuropathy. Int J Mol Sci 20, 2019

100. Zhang H, Li Y, de Carvalho-Barbosa M, Kavelaars A, Heijnen CJ, Albrecht PJ, Dougherty PM: Dorsal Root Ganglion Infiltration by Macrophages Contributes to Paclitaxel Chemotherapy-Induced Peripheral Neuropathy. J Pain 17:775–86, 2016

101. Zhang XC, Levy D: Modulation of meningeal nociceptors mechanosensitivity by peripheral proteinase-activated receptor-2: the role of mast cells. Cephalalgia 28:276–84, 2008

